# MicroRNA-205 affects mouse granulosa cell apoptosis and estradiol synthesis by targeting CREB1

**DOI:** 10.1101/301523

**Authors:** Pengju Zhang, Jun Wang, Hongyan Lang, Weixia Wang, Xiaohui Liu, Haiyan Liu, Chengcheng Tan, Xintao Li, Yumin Zhao, Xinghong Wu

**Affiliations:** Institute of Animal Sciences, Jilin Academy of Agricultural Sciences, #1363 Shengtai Street, Changchun 130124, Jilin Province, PR China; College of Animal Science and Technology, Jilin Agricultural University, 2888 Xincheng Street, Changchun 130118, Jilin Province, PR China

**Keywords:** Follicular atresia, miR-205, Apoptosis, Estradiol synthesis, CREB1

## Abstract

MicroRNAs-205 (miR-205), were reportedly to be involved in various physiological and pathological processes, but its biological function in follicular atresia remain unknown. In this study, we investigated the expression of miR-205 in mouse granulosa cells (mGCs), and explored its functions in primary mGCs using a serial of *in vitro* experiments. The result of qRT-PCR demonstrated that miR-205 expression was significantly increased in early atretic follicles (EAF), and progressively atretic follicles (PAF) compared to healthy follicles (HF). Our results also revealed that overexpression of miR-205 in mGCs significantly promoted apoptosis, caspas-3/9 activities, and inhibited estrogen E2 release, and cytochrome P450 family 19 subfamily A polypeptide 1 (CYP19A1, a key gene in E2 production) expression. Bioinformatics and luciferase reporter assays revealed that the gene of cyclic AMP response element (CRE)-binding protein 1 (CREB1) was a potential target of miR-205. qRT-PCR and western blot assays revealed that overexpression of miR-205 inhibited the expression of CREB1 in mGCs. Importantly, CREB1 upregulation partially rescued the effects of miR-205 on apoptosis, caspase-3/9 activities, E2 production and CYP19A1 expression in mGCs. Our results indicate that miR-205 may play an important role in ovarian follicular development and provide new insights into follicular atresia.

## INTRODUCTION

A large number of follicles form in the ovaries of humans and domestic animals during puberty, and each follicle contains an oocyte surrounded by several layers of granulosa cells (GCs)(1). It is well known that most ovarian follicles (> 99%) undergo a degenerative process known as atresia at different stages of follicular development, and only a limited number of follicles ovulate(<1%)(2,3). Therefore, it is crucial important to prevent follicular atresia for the recruitment of follicles and subsequent ovulation. Ovarian GC apoptosis was reportedly to play a major role in the fate of follicles at all stages of follicular development, and was regarded as the main cause of follicular atresia (4,5). It has been demonstrated that high incidence of GC apoptosis caused empty follicles increased, fewer oocyte retrieval, and oocyte quality reduced (6). To elucidate the mechanism underlying follicular atresia, it is an urgent need to investigate the effect GC apoptosis.

Estrogens are required for female reproduction, and are synthesized from ovarian androgen in GC by a rate limiting enzyme aromatase (7,8). Aromatase enzyme is encoded by the gene factor cytochrome P450 family 19 subfamily A polypeptide 1 (CYP19A1)(9). E2, a major product of estrogen, has been shown to be involved in follicular atresia by stimulating GC proliferation and protecting GC apoptosis (10,11).

In recent years, microRNAs (miRNAs) have been detected in the ovarian tissues and have been shown to be involved in ovarian development, oocyte maturation, follicular atresia, and embryo development (12,13). Several miRNAs has been identified to play crucial in follicular atresia (12). For example, Liu *et al* reported that miR-1275 promoted apoptosis and estradiol synthesis by impairing LRH-1/CYP19A1 axis in pGCs(14). Yao *et al* showed that miR-181b-induced SMAD7 downregulation controls GCs apoptosis through TGF-β signaling by interacting with the TGFBR1 promoter (15). Xu *et al* found that miR-145 protects GCs against oxidative stress-induced apoptosis by targeting KLF4 (16). These results suggesting that ovarian functional studies related to miRNA regulation become a study hot.

MicroRNA-205 (miR-205), as a highly conserved microRNA, has been demonstrated to play key mediatory roles in multiple cellular processes such as cell survival, proliferation, apoptosis, angiogenesis, and embryonic development (17). It has been shown that miR-205 might be involved in oocyte maturation and embryo development in pigs and bovine (18,19) However, the regulatory role and molecular mechanisms of miR-205 in GC apoptosis during follicle atresia remain unclear. In this study, we investigated the involvement of miR-205 in mouse GCs (mGCs) apoptosis and estradiol synthesis. We also investigated the regulatory mechanism of miR-205 in mGCs by a serial of molecular and cell experiments *in vitro*.

## RESULTS

### miR-205 expression level was increased during follicular atresia

The expression levels of miR-205 in healthy follicles (HF), early atretic follicles (EAF), and progressively atretic follicles (PAF) were detected by qRT-PCR. The data revealed that miR-205 expression levels were significantly increased in EAF and PAF groups compared with HF (both *P*<0.05, Fig.1), suggesting that miR-205 plays a crucial role in follicular atresia.

**Fig.1.**
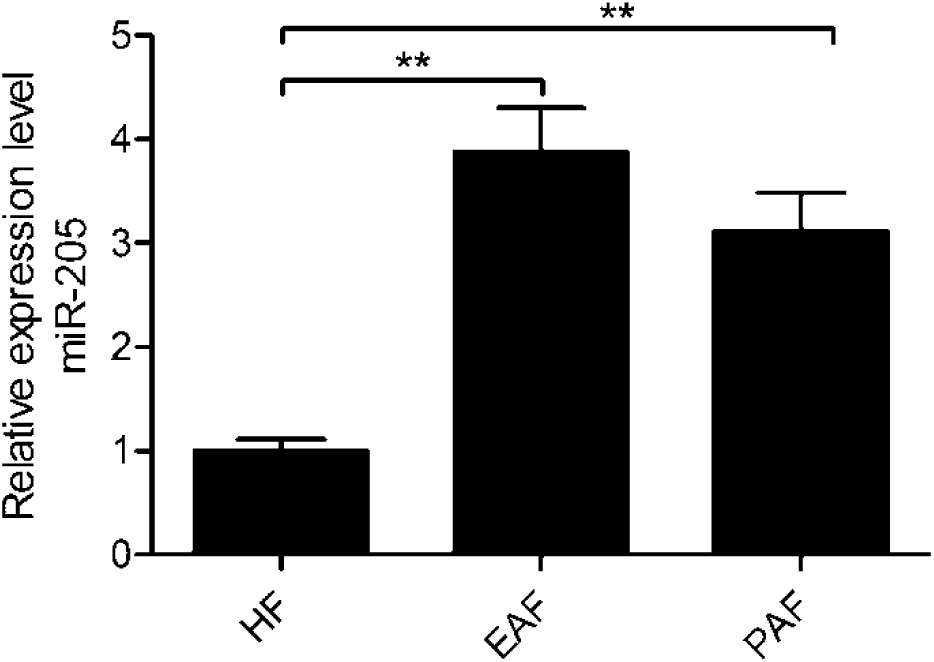
The expression of miR-205 in mouse ovarian granulosa cells. The relative expression levels of miR-205 in healthy follicles (HF), early atretic follicles (EAF), and progressively atretic follicles (PAF) were determined using qRT-PCR. U6 was used as a loading control to normalize expression levels. ^**^*P*<0.05, ^**^*P*<0.01.

### Overexpression of miR-205 promoted mGC cell apoptosis and caspas-3/9 activities

To evaluate miR-205 effect on mGC apoptosis, the mGCs were transfected with miR-205 mimics or miR-NC. The results showed that mGCs transfected with miR-205 mimics significantly enhanced miR-205 expression compared with mGCs transfected with miR-NC (Fig.2A). Next, we investigated the potential effect of miR-205 on mGCs apoptosis. Flow cytometric analysis revealed that overexpression of miR-205 significantly induced mGC apoptosis (Fig.2B). Moreover, we found that overexpression of miR-205 significantly promoted caspase-3/9 activities (Fig.2C, D), increased Bax protein expression (a pro-apoptotic factor), and decreased Bcl-2(a anti-apoptotic factor) in mGCs (Fig. 2E).Collectively, these data clearly show that miR-205 can inhibit mGC apoptosis.

**Fig.2.**
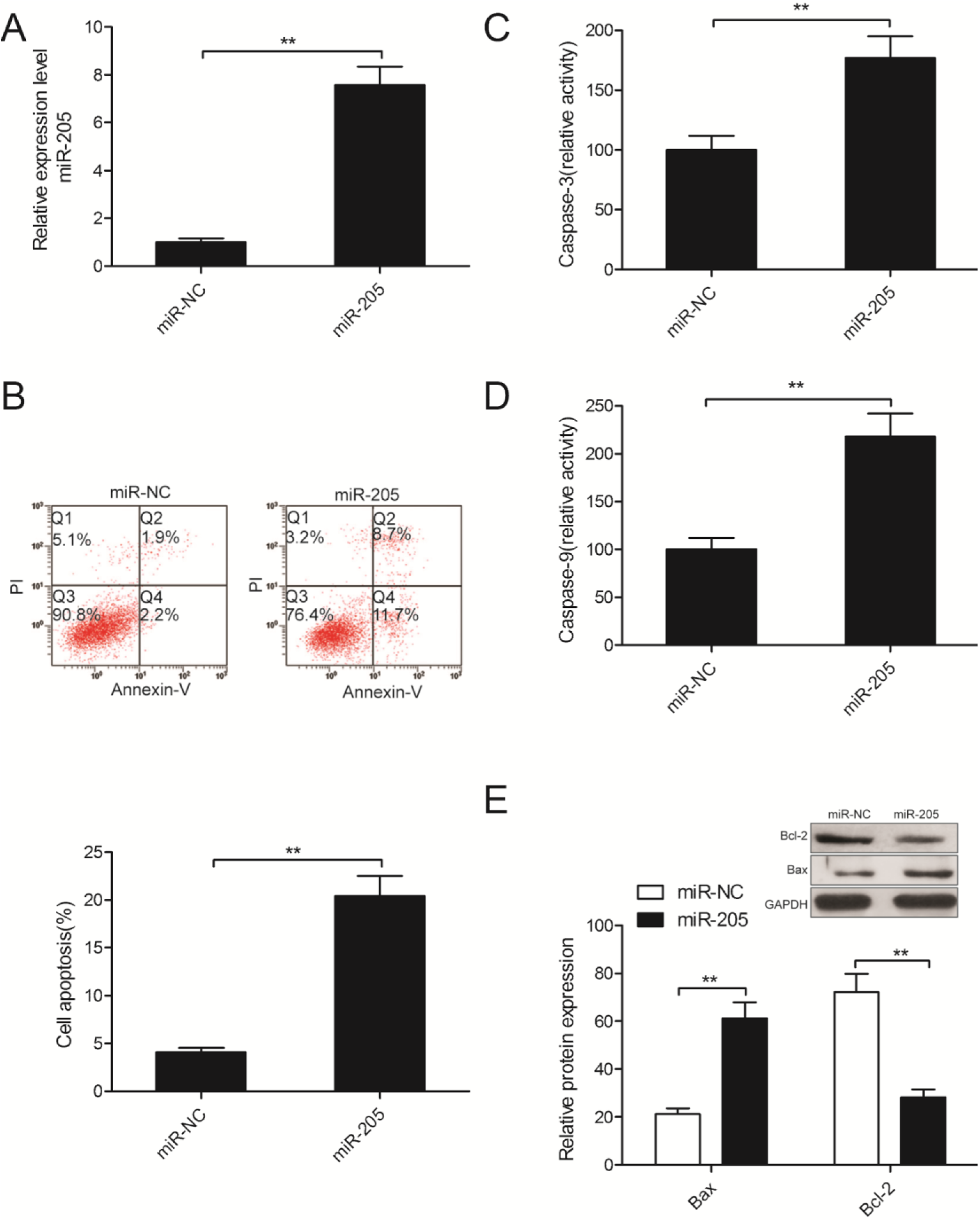
Overexpression of miR-205 promoted mGC cell apoptosis and caspas-3/9 activities. (A) The relative expression levels of miR-205 were detected in mGCs transfected with miR-205 mimics or miR-NC. U6 was used as a loading control to normalize expression levels. (B) Cell apoptosis was determined in mGCs transfected with miR-205 mimics or miR-NC by flow cytometric analysis. (C,D) Caspas-3/9 activities were detected in mGCs transfected with miR-205 mimics or miR-NC. (E) Bax and Bcl-2 protein expression were detected in mGCs transfected with miR-205 mimics or miR-NC by western blot. GAPDH was used as a loading control to normalize expression levels. ^*^*P*<0.05, ^**^*P*<0.01.

### miR-205 suppresses E2 synthesis in mGCs

Follicular atresia was closely associated with correlates with estrogen E2 concentrations in follicular fluid (14, 20). To further investigate the effect of miR-205 on estrogen, the concentration of E2 in the medium of mGCs transfected with miR-205 mimics was measured using a RIA kit. Our results revealed that overexpression of miR-205 significantly decreased E2 release in mGCs (Fig.3A). CYP19A1, a key enzyme in E2 synthesis, was reportedly to play a key role in during atresia follicles (14, 21) Thus, we detected CYP19A1 expression on mRNA and protein levels in mGCs transfected with miR-205 mimics by qRT-PCR and western blot, respectively. Our results showed that overexpression of miR-205 significantly decreased the mRNA (Fig.3B) and protein (Fig.3C) expression of CYP19A1 in mGCs. These results implied that miR-205 can regulate E2 synthesis in mGCs by regulating CYP19A1 expression, leading to mGC apoptosis and follicular atresia.

**Fig.3.**
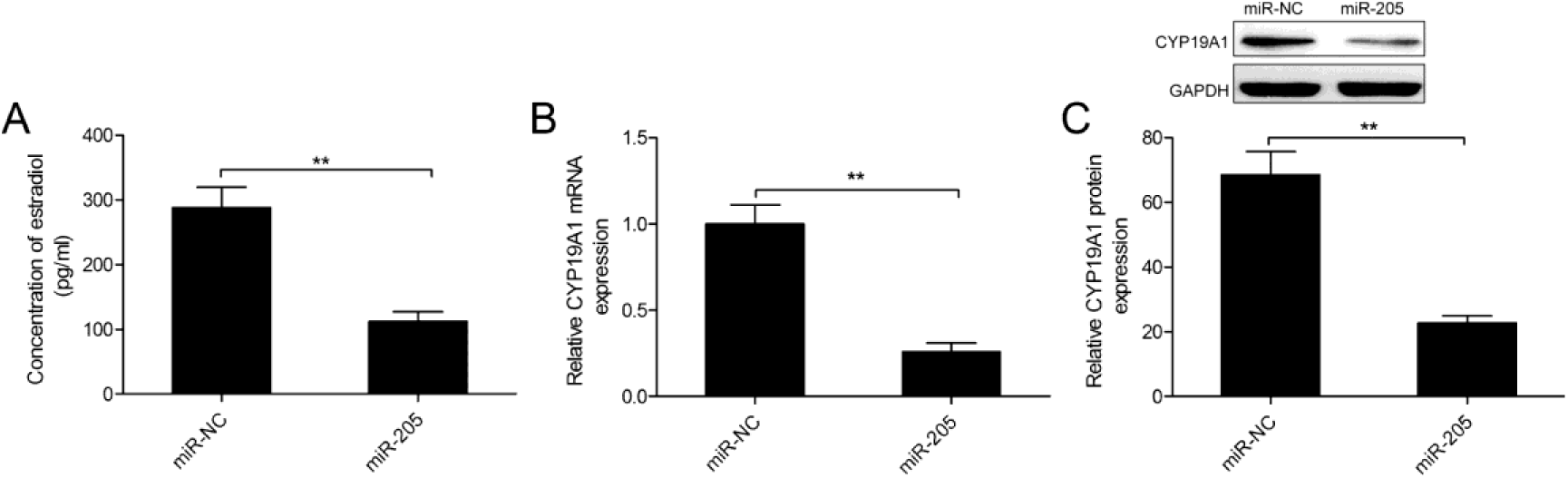
miR-205 suppresses E2 synthesis in mGCs. (A) The concentration of E2 was measured in mGCs transfected with miR-205 mimic or miR-NC. (B,C) The CYP19A1 expression on mRNA and protein levels was detected in mGCs transfected with miR-205 mimics or miR-NC by qRT-PCR and western blot, respectively. GAPDH was used as a loading control to normalize expression levels. ^*^*P*<0.05, ^**^*P*<0.01.

### CREB1 is a direct target of miR-205 in mGCs

To determine the mechanism of action of miR-205 in mGCs cells, we performed a miRNA target search using TargetScan and miRWalk. We selected CREB1 as a potential downstream target gene of miR-205 based on CREB1 function in ovarian GC (22,23). As shown in Fig.4A, a highly conserved putative miR-205 recognition sequence binds to the 3′-UTR of *CREB1* at positions 2036–2043, suggesting that this gene may be a target of miR-205. To further confirm targeting of CREB1 by miR-205, luciferase activity assay was performed in HEK293 cells. Our results demonstrate that miR-205 obviously inhibited the luciferase activity of the wild-type (WT) of CREB1 −3’UTR but not that of the mutant-type (MT) 3’ UTR of CREB1 (Fig.4B). Subsequently, we examined the CREB1 expression on mRNA and protein levels in mGCs transfected with miR-205 or miR-NC by qRT-PCR and Western blot, respectively. We found that overexpression of miR-205 in mGCs significantly downregulated CREB1 expression both mRNA and protein levels (Fig.4C, D). These results suggested that CREB1 might be a target of miR-205 in mGCs.

**Fig.4.**
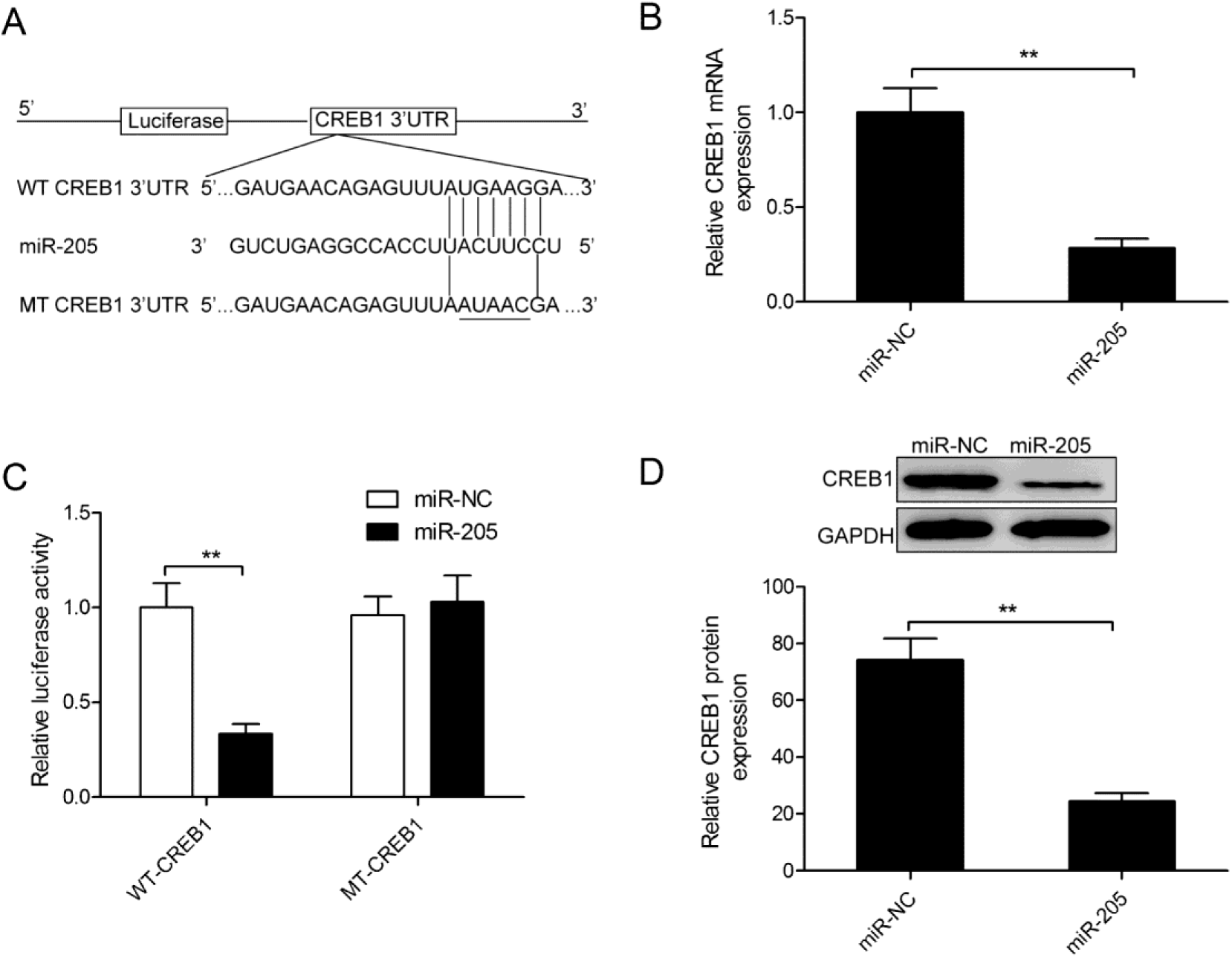
CREB1 is a direct target of miR-205 in mGCs. (A) Sequence alignment of wild-type (WT) and mutated (MT) putative miR-205-binding sites in the 3′-UTR of CREB1. (B) Luciferase reporter assays were performed using HEY293 cells co-transfected with the miR-205 mimics or miR-NC and *CREB1*-WT-3′-UTR or CREB1-MT-3′-UTR reporter plasmid.(C,D) The CREB1 expression on mRNA and protein levels was detected in mGCs transfected with miR-205 mimics or miR-NC by qRT-PCR and western blot, respectively. GAPDH was used as a loading control to normalize expression levels. ^*^*P*<0.05, ^**^*P*<0.01.

### Overexpression of CREB1 partially attenuated the effects of miR-205 in mGCs

To validate whether miR-205 affects mGC apoptosis and estradiol synthesis via targeting CREB1, mGCs with miR-205 mimics or miR-NC were transfected pcDNA3.1-CREB1 plasmid (without the 3′-UTR).It was found that the forced expression of the CREB1 could recover CREB1 expression on mRNA levels and protein level in mGCs transfected with miR-205 (Fig.5A,B).In addition, restoration of CREB1 expression in mGCs partially reversed the effects of the miR-205 on Bax, Bcl-2, and CYP19A1 expression, cell apoptosis, caspase-3/9 activity, and estradiol synthesis (Fig.5B-F). These data indicate that miR-205 affects mGC apoptosis and estradiol synthesis, in part by suppressing *CREB1* expression.

**Fig.5.**
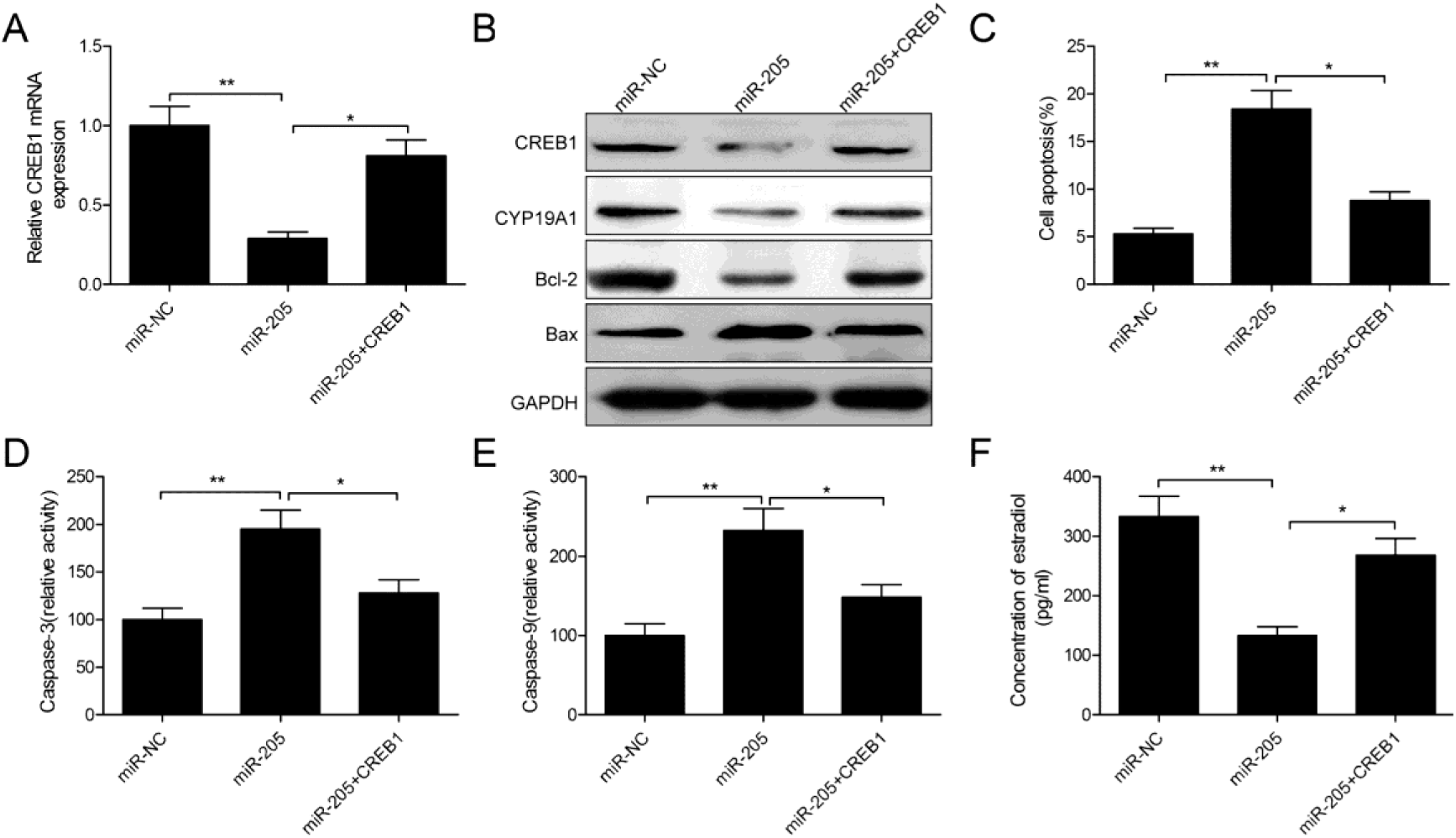
Overexpression of CREB1 partially attenuated the effects of miR-205 in mGCs. (A) *CREB1* mRNA expression was determined in mGCs transfected with miR-205 mimics or miR-NC plasmid and with/without the overexpressing CREB1 plasmid (without3’UTR). GAPDH was used as a loading control to normalize expression levels. (B) CREB1 Bax, Bcl-2 and CYP19A1 proteins expression were determined in mGCs transfected with miR-205 or miR-NC plasmid and with/without the overexpressing CREB1 plasmid (without3’UTR). GAPDH was used as a loading control to normalize expression levels. GAPDH was used as a loading control to normalize expression levels. (C-F) The effect of miR-205 on cell apoptosis,caspas-3/9 activities, and E2 concentration in mGCs were reversed under the condition of overexpression of CREB1. ^*^*P*<0.05, ^**^*P*<0.01.

## DISCUSSION

MicroRNAs (miRNAs) are small (18–25 nucleotide in length), single stranded, noncoding RNAs which inhibit the expression of target genes by binding to 3’-untranslated region (UTR) of their target mRNA transcripts, leading to mRNA destruction or translational inhibition (24). It has been gradually clear that miRNAs are involved in regulating various ovarian functions, including GC proliferation, GC apoptosis, ovulation, corpus luteum development, as well as follicular atresia(13,25). In the present study, we investigate the biological role of miR-205 in mGC progression. Our results demonstrated that miR-205 expression was upregulated in EAF and PAF. Overexpression of miR-205 in mGCs significantly promoted apoptosis, and caspase-3/9 activities, and decreased E2 concentration and CYP19A1 expression, suggesting that miR-205 could control ovarian GCs apoptosis and steroid synthesis. To our knowledge, we for the first time demonstrated that miR-205 regulated apoptosis and steroid synthesis in mGCs.

Growing evidences has shown that miR-205 expression was upregulated in ovarian cancer tissues, and functioned as oncogene in tumor initiation and development of ovarian cancer by regulating multiple target genes (26–29). Moreover, overexpression of miR-205 could stimulate formation and adhesion of extra-embryonic endoderm cells from pluripotent embryonic stem cells or trigger pluripotent cell progenitors and modulate spermatogenesis (30). Importantly, miR-205 was reported to be involved in oocyte maturation and embryo development in pigs and bovine (18,19). These studies implied that miR-205 might play crucial mediatory roles in ovarian disease, oocyte maturation and embryo development. However, the regulatory role and molecular mechanisms of miR-205 in GC apoptosis during follicle atresia were unknown. Our results showed that miR-205 expression was upregualted in EAF and PAF, suggesting miR-205 was involved in follicular atresia progression. Function assays shown that overexpression of miR-205 promoted mGC apoptosis and caspase-3/9 activity, and decreased steroid synthesis. These results suggested that miR-205 could affect follicular atresia progression.

To further investigate the molecular mechanisms underlying the influence of miR-205 on mGC apoptosis and steroid synthesis, we used two algorithms (TargetScan and miRWalk) to identify its putative protein-coding gene targets, particularly those known to affect GC apoptosis and steroid synthesis. From this, cAMP responsive element binding protein 1 (*CREB1*) was selected, given its close association with GC apoptosis and steroid synthesis (22,23). Activated CREB1 can regulate the expression of multiple downstream genes, such as those encoding proteins involved in apoptosis (BCL-2), the cell cycle (cyclinA1, B1, and D2), and signal transduction (activating transcription factor 3 and NF-κB), and steroid synthesis (CYP19A1)(31–33). CREB1 had been identified as a target of miR-205 in colorectal cancer and neuroblastoma (34,35). However, it is unclear whether CREB1 can serve as a target of miR-205 in mGC. Here, we further confirmed that *CREB1* was a miR-205 target by luciferase reporter assay. qRT-PCR and western blotting also revealed that expression of miR-205 mimic in mGC cells significantly inhibited CREB1 expression, showing that miR-205 targets *CREB1* in mGC. Of note, CREB1 overexpression reversed effect on cell apoptosis, caspase-3/9activities, and steroid synthesis in mGC mediated by miR-205 overexpression. These findings suggested that miR-205 control ovarian GCs apoptosis and steroid synthesis, at least in part, by repressing CREB1.

In summary, the present investigation revealed that miR-205 affects ovarian GCs apoptosis and steroid synthesis, at least to a certain extent by inhibiting CREB1. This findings provided an improved understanding of molecular mechanisms of miR-205 regulating ovarian follicular atresia, suggesting miR-205 may be a potentially target for follicular atresia.

## MATERIALS AND METHODS

### Animals and follicle isolation

C57BL/6J female mice were obtained from the Experimental Animal Center of Jilin University (Changchun, China), and were housed under a 12 h light/12 h dark schedule and provided normal food and water ad libitum. All animal experiments were conducted according to the guidelines for the welfare and use of animal in cancer research (ad hoc committee of the National Cancer Research Institute, UK), and were approved by the Ethics Committee of Jilin Academy of Agricultural Sciences(Changchun, Jilin).

Follicles were isolated from ovaries of mature C57BL/6J female mice, and then were divided into healthy follicles (HF), early atretic (EAF), and progressively atretic (PAF) follicles according to their morphologic characteristics.

### Quantitative real-time polymerase chain reaction (qRT-PCR)

Total RNA were extracted from follicles or cultured mGCs using a TRIZOL regent kit (Invitrogen, Carlsbad, CA, USA) according to the manufacturer’s instructions. The purity of RNA was determined by a NanoDrop 2000 UV-Vis-spectrophotometer (NanoDrop Technologies, Wilmington, DE, USA) at 260/280. To detect miR-205, total RNA was reversely transcribed into cDNA using TaqMan^®^ MicroRNA Reverse Transcription Kit (Applied Biosystems, Foster City, CA, USA), and was quantified using miScript SYBR Green PCR Kit (QIAGEN, Hilden, Germany) on an ABI 7900 Real-Time PCR System (Applied Biosystems). To quantify *CREB1* and *CYP19A1* mRNA levels, total RNA was reversely transcribed using PrimeScript RT Reagent Kit (Takara, Dalian, China) and then was quantified with Real-time PCR Mixture Reagent (Takara) on an ABI 7900 Real-Time PCR System. The primer sequences are listed in Table 1, and the detail conditions were as follows: 95°C for 5 min, followed by 40 cycles of 95°C for 20 s and 58°C for 1 min. Relative miR-205 and *CREB1* mRNA expression was quantified using the 2^−ΔΔ^Ct method to *U6* and *GAPDH* levels, respectively.

**Table 1.**
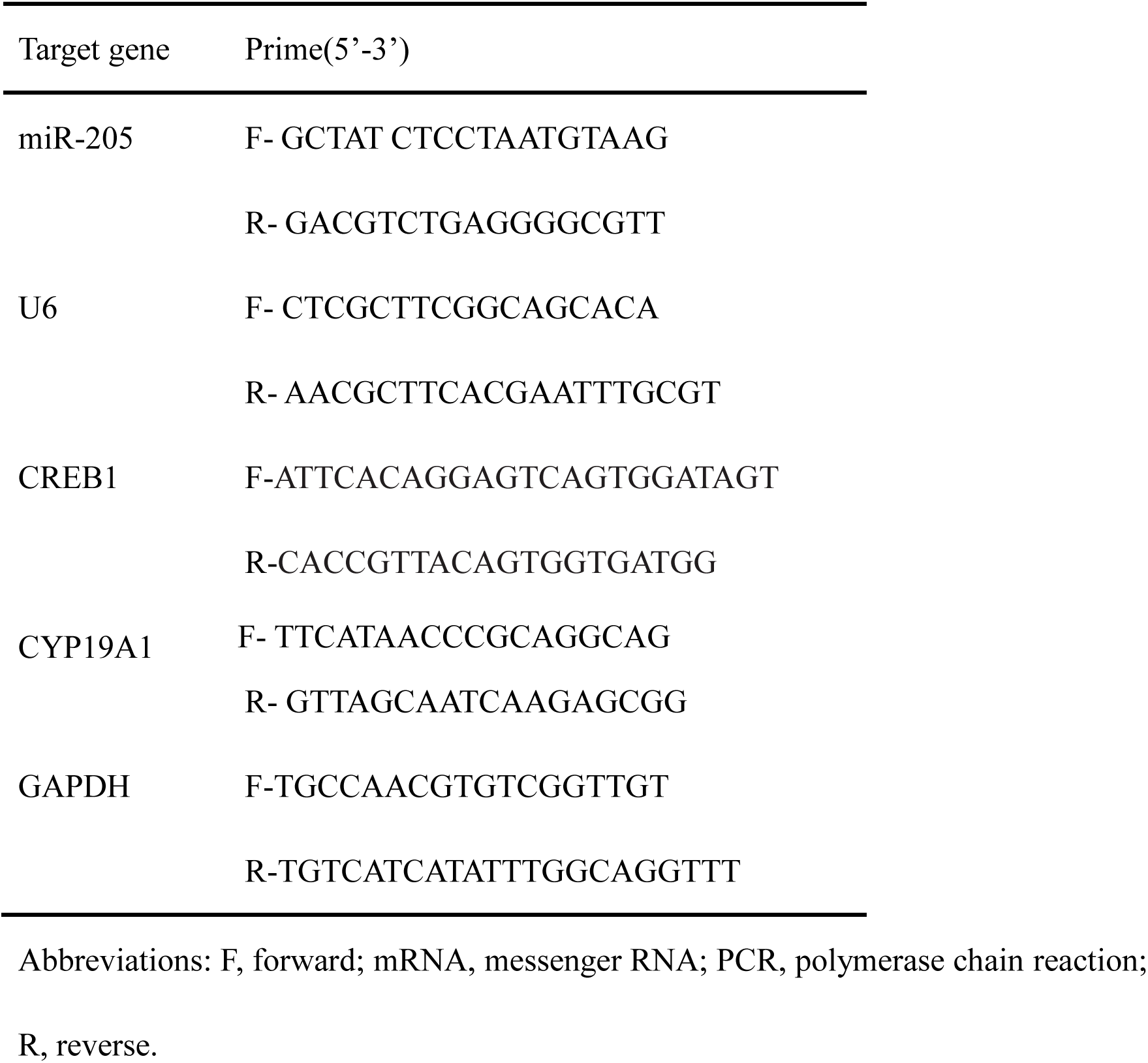
Primers for qRT-PCR analysis.

### Isolation and culture of primary mouse granulosa cells (mGCs)

Mouse primary GCs (mGCs) were isolated from the ovaries of 21-day-old immature C57BL/6 mice by the follicular puncture method as described by Zhang et al (36). mGCs were cultured in DMEM/F12 (Sigma, USA) containing with 10 % heat-inactivated fetal bovine serum (FBS, Gibco BRL), penicillin 100 U/ml) and streptomycin (100 U/ml) (Life Technologies, CA) at 37 °C in a humidified atmosphere containing 5% CO2. The media were changed every 2 day. The second or third passage mGCs were selected for subsequent experiments. HEK293 cells were brought from American Type Culture Collection (ATCC, VA, USA), and were cultured in DMEM containing 10% FBS and 1% penicillin/streptomycin in 5% CO2 at 37 °C.

### Transient transfection

miR-205 mimic and corresponding negative mimic (miR-NC) were chemically synthesized from GenePharma Co., Ltd. (Shanghai, China).The *CREB1* overexpression plasmid (pCDNA3.1-*CREB1*) was granted from Dr Xue Wang (Fudan University). mGCs in logarithmic phase were transfected with above-mentioned molecular production using Lipofectamine 2000 (Invitrogen) according to the manufacturer’s instructions.

### Flow cytometric analysis of apoptosis

The transfected mGCs were cultured for 48 h, and then were harvested. The apoptotic cells were detected using an ApoScreen Annexin V Apoptosis Kit (Southern Biotech, Birmingham, AL, USA) following the manufacturer’s instructions. The stained cells were examined using fluorescence-activated cell sorting (FACS) using a flow cytometer (BD Biosciences, Franklin Lakes, NJ, USA) with a single 488 nm laser excitation source. Apoptosis was analyzed using CellQuest software (BD Biosciences).

### Caspase-3/-9 activity assay

Next, caspase-3/-9 activities were determined using a Caspase-3/-9 Activity Assay Kit (Beyotime, Beijing, China) according to the manufacturer’s protocol. Absorbance at a wavelength of 405 nm was measured in a microplate spectrophotometer (Thermo Labsystems, Vantaa, Finland).

### Radioimmunoassay (RIA)

Transfected mGCs were cultured in FBS-free DMEM/F12 (phenol red-free) medium for 24 h. The medium was then collected by centrifugation at 3000 ×g for 5 min. Estradiol E2 concentrations were determined at the First of Hospital of Jilin University (Changchun, China) using radioimmunoassay (RIA) kits (Beckman) according to the manufacturer’s protocol. The sensitivities of the E2 assays are 2 pg/mL.

### Western blot

The cultured cells were collected and lysed using RIPA buffer (Sigma-Aldrich) according to the protocol of manufacture. After total protein quantification by BCA kit (Sigma-Aldrich), an equal quantity 30μg protein each well was separated by 10% sodium dodecyl sulfate polyacrylamide gel electrophoresis (SDS–PAGE), and then electrophoretic transferred onto PVDF membrane (Millipore, MA, USA). After blocking with 5% non-fat dry milk in Tris-buffered saline with Tween-20 (TBS-T), the membranes were probed with primary anti-CREB1 antibody (1:1000 dilution; Santa Cruz Biotechnology Inc., Santa Cruz, CA, USA), anti-Bax(1:1000 dilution; Santa Cruz Biotechnology Inc), anti-Bcl-2(1:1000 dilution; Santa Cruz Biotechnology Inc), anti-CYP19A1(1:1000 dilution; Santa Cruz Biotechnology Inc) and anti-GAPDH antibody (1:3000 dilution; Santa Cruz Biotechnology Inc) at 4 °C overnight, followed by incubation with horseradish peroxidase (HRP)-conjugated anti-mouse IgG secondary antibody (1:5000 dilution; Santa Cruz Biotechnology Inc) at 37 °C for 2 h. GAPDH was used as an internal control. The protein bands were visualized using an ECL detection reagent on Bio-Rad ChemiDoc MP (Bio-Rad,USA). The relative protein intensities were analyzed using Gel-pro Analyzer^®^ software (Media Cybernetics, Rockville, MD, USA).

### Luciferase reporter assay

The binding sites between miR-205 and CREB1 were predicted using targetscan (http://www.targetscan.org/),andmiRWalk(http://zmf.umm.uni-heidelberg.de/apps/zmf/mirwalk/index.html). Mouse *CREB1* 3′-untranslated region (3′UTR)′sequences containing putative or mutated miR-205 binding site, were chemically synthesized and cloned into the downstream of psiCHECK-2 vector (Promega, Madison, WI, USA). These constructs were termed CREB1-WT-3′-UTR and CREB1-MT-3’-UTR, respectively. HEK293 cells were seeded into 24-well plates and grown to 70 – 80% confluent. Then cells were co-transfected with the CREB1-WT-3’-UTR or *CREB1*-MT-3′-UTR reporter plasmid (100 ng) and the miR-205 mimic or miR-NC (100 nM) using Lipofectamine 2000 (Invitrogen). 48 h after transfection, firefly and renilla luciferase activities in cell lysates were measured using the Dual-Luciferase Reporter System (Promega). The relative luciferase activity was standardized with renilla luciferase activity.

### Statistical analysis

Qualitative data obtained from the at least three times replicate experiments and presented as mean ±standard deviations(SD). Statistical analysis was conducted using SPSS 18.0 (SPSS Inc., Chicago, IL, USA) and GraphPad Prism 5 (GraphPad Software, Inc., San Diego, CA, USA). The *t*-test was used to compare between two groups, and one-way analysis of variance (ANOVA) was adopted for comparisons more than two groups. A *P* value<0.05 was considered statistically significant.

## Acknowledgements

We thank Prof. Na Li, Department of Obstetrics and Gynecology, The First Hospital of Jilin University for RIA analysis.

## Author contributions

Pengju Zhang, Xintao Li, Yumin Zhao and Xinghong Wu conceived and designed the study; Pengju Zhang, Jun Wang, Hongyan Lang, Weixia Wang, Xiaohui Liu performed the experiments; Haiyan Liu analyzed the data, Chengcheng Tan discussed results and advised during the completion of the study; Pengju Zhang, Xintao Li, Yumin Zhao and Xinghong Wu wrote the paper.

